# Camera-Based Bi-Axial Measurement of Weak Forces Generated by Freely-Moving Plant Organs

**DOI:** 10.1101/2025.05.29.656568

**Authors:** Amir Ohad, Yasmine Meroz

## Abstract

Growing plants are remarkable at negotiating obstacles in their unstructured and changing environments. Measuring the mechanical interactions of growing plants with surrounding objects is a critical step towards deciphering thigmotropic responses underpinning complex growth strategies. Yet, available force-measurement systems have limited capacity to capture weak (sub-mN) forces in freely moving plant organs - such as the forces applied by a growing shoot pushing at an obstacle. We developed a measurement system based on the deflection of a pendulum by a freely moving shoot. Crucially, unlike many force-measurement systems, the organ is not tethered to the device. Moreover, force is measured along two axes, as opposed to one axis in commonly used methods such as cantilevers. Orthogonal cameras track the 3D position of the rod and shoot, yielding the rod deflection angle and, using a mechanical torque equilibrium equation, allowing to extract the force applied by the plant over time. This system is relevant for measuring weak forces in macro-sized systems (such as growth or turgor pressures), and the force detection range can be tuned by altering rod mass and length. We demonstrate the system with bean (*Phaseolus vulgaris*) shoots, measuring the forces they apply on a candidate support during inherent circumnutation movements, prior to twining. Such measurements lay the foundations for deciphering how climbing plants assess whether to twine or not - an open question since Darwin’s first observations.

## Introduction

As plants grow and forage for nutrients and light, they come into contact with surrounding objects, and engage in mechanical interactions with their environment (Darwin, 1875; Gianoli, 2015; Moulia et al., 2022). Such interactions lead to complex growth patterns (Tedone et al., 2020; Goriely, 2017; Goriely & Neukirch, 2006; Chelakkot & Mahadevan, 2017), as showcased in observations of waving and coiling of roots grown on inclined substrates (Okada & Shimura, 1990; Thompson & Holbrook, 2004; Zhang et al., 2022; Porat et al., 2024), and ultimately determine a plant’s capacity to negotiate its environment. This idea is particularly salient in the case of thigmomorphogenesis (Telewski & Jaffe, 1986; Jaffe, 1973; Biddington, 1986; Jaffe et al., 2002) and thigmotropism (Braam, 2004; Forterre et al., 2005; Takahashi & Jaffe, 1990): plants’ active growth responses to mechanical stimuli. These growth responses enable plants to maintain structural integrity in the face of environmental threats such as wind (Telewski & Jaffe, 1986; Moulia et al., 2015) and herbivore damage (González-Teuber & Gianoli, 2007).

Clearly, measuring mechanical interactions between plants and their environment is a critical step towards characterizing and eventually deciphering the complex growth strategies of plants. Yet, researchers have faced challenges in measuring these interactions in non-tethered plant organs, where current efforts have largely focused on root-soil systems (Kolb et al., 2017, 2022; Quiros et al., 2022; Koren et al., 2024), where force measurements are typically conducted by clamping or restricting the organ. However, techniques for studying unrestricted plant movements are still lacking. An important case in point is the dynamic twining response of climbing plants encountering a support (Darwin, 1875; von Sachs, 1887; Silk, 1989; Gianoli, 2015), during inherent sweeping semi-oscillatory movements termed circumnutations (CN) (Darwin, 1875; von Sachs, 1887; Stolarz, 2009; Gianoli, 2015; Bai et al., 2024)(Rivière et al., 2025), a classic, and experimentally tractable example of thigmomorphogenesis.

Though multiple studies measure forces exerted by attachments of climbing plants on supporting structures (Jaffe, 1970a; Silk & Hubbard, 1991; Matista & Silk, 1997; Silk & Holbrook, 2005; Steinbrecher et al., 2010; Bauer et al., 2010), and recent studies measure the forces exerted on a support by coiling tendrils (Klimm et al., 2023, 2024), very little work focuses on the initiation of twining ((Jaffe, 1970a; Goriely & Neukirch, 2006)), and there are no quantitative measurements of the resistance exerted by newly encountered supports on twiners prior to twining.

Such measurements could shed light the on twining initiation of stem twiners and other climbing plants. The challenge in measuring such forces, rises from a broader difficulty in measuring low (sub-mN) forces of freely moving organs.

The forces generated by these motions are expected to be of low magnitude (even a light breeze will stir a thin stem), requiring sensitive measurements. Though numerous systems have been developed to measure low forces (see examples in Table 1), most are designed to measure forces applied by millimeter to micrometer (or smaller) size organisms. Consequently, the measurement devices and setups are of comparable dimensions, which presents a challenge for measuring small forces in large (centimeter-scale and higher) organisms, such as in climbing plants (Klimm et al., 2023; Jaffe, 1970b). Replicating existing measurements systems at a larger scale is either not feasible, or fundamentally not suitable for force measurements of freely moving circumnutations altogether. For example, many existing systems are based on measuring deflection of elastic cantilevers (Kamimura & Takahashi, 1981; Kishino & Yanagida, 1988; Beaussart et al., 2014; Backholm & Bäumchen, 2019; Poppele & Hozalski, 2003), yielding a 1D projection of the applied force parallel to the measurement axis of the device (Fig. 1A), and typically requiring tethering of the sample to secure the direction of the applied force. While this method has been successfully adapted for plant roots (Quiros et al., 2022), in the case of a circumnutating plant organ, tethering significantly alters the inherent stem dynamics, and therefore the force generation itself.

**Table 1.** Table of example previous force measurement setups in living systems. Methods of measurement labeled by broad type, sample refers to the type of biological system involved in the force measurement. Sample sizes are the lengths of samples presented for measurement, or estimation of possible sizes of measured samples in the setup. Force generation labels the form in which the forces are generated, or how they are used (as in elastic modulus measurements). The measurement axes describe the axes in which force can be measured simultaneously with the given setup, and the force range gives an estimation of the possible forces available for measurement within the setup limits. PIV - particle image velocimetry. AFM - atomic force microscopy.

| Reference | Method | Sample | Sample size (m) | Force generation | Measurement Axes | Force range (N) |
| --- | --- | --- | --- | --- | --- | --- |
| Kamimura & Takahashi (1981) | Cantilever | Dynein arm | $10^{-7} - 10^{-4}$ | Sliding | 1 | $10^{-12} - 10^{-10}$ |
| Kishino & Yanagida (1988) | Cantilever | Actin | $10^{-6} - 10^{-4}$ | Tensile | 1 | $10^{-12} - 10^{-10}$ |
| Autumn et al. (2000) | AFM cantilever | Gecko seta | $10^{-6} - 10^{-4}$ | Adhesion | 2 | $10^{-7} - 10^{-4}$ |
| Poppele & Hozalski (2003) | Cantilever | Biofilms | $10^{-6} - 10^{-3}$ | Adhesion | 1 | $10^{-7} - 10^{-5}$ |
| Doll et al. (2009) | Strain gauge | Nematodes | $10^{-6} - 10^{-4}$ | Propulsion | 2 | $10^{-6} - 10^{-4}$ |
| Sznitman et al. (2010) | PIV | Nematodes | $> 10^{-5}$ | Propulsion | 2 | $> 10^{-9}$ |
| Rabets et al. (2014) | Cantilever | Nematodes | $< 10^{-2}$ | Drag forces | 2 | $10^{-6} - 10^{-3}$ |
| Beaussart et al. (2014) | Force microscopy | Microbial | $10^{-9} - 10^{-6}$ | Adhesion | 1 | $10^{-11} - 10^{-8}$ |
| Backholm & Bäumchen (2019) | Cantilever | Multiple | $10^{-6} - 10^{-3}$ | Multiple | 1 | $10^{-11} - 10^{-3}$ |
| Das et al. (2022) | Force transducer | Neurons | $> 10^{-3}$ | Elastic | 1 | $> 10^{-3}$ |
| Filc et al. (2023) | Force transducer | Wings | $10^{-3} - 10^{-1}$ | Elastic | 1 | $10^{-4} - 10^{-2}$ |
| Klimm et al. (2023) | Force transducer | Tendrils | $10^{-2} - 10^0$ | Tension | 3 | $10^{-3} - 10^0$ |

**Fig. 1.**
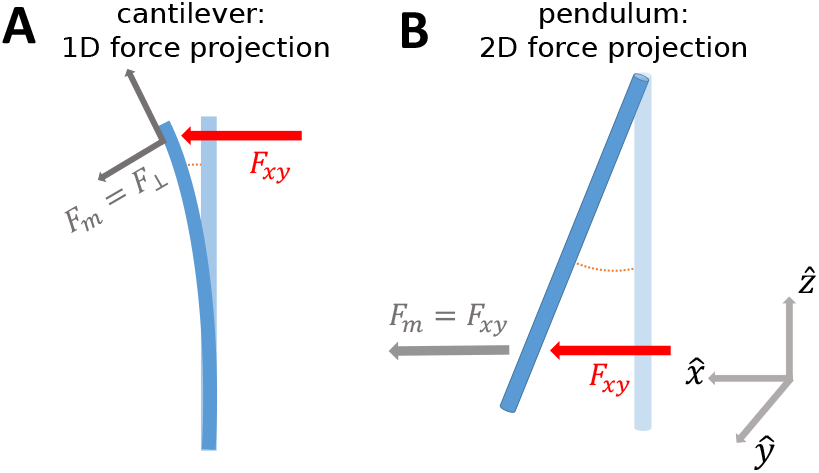
Comparison of measured force projections of common cantilever-based force measurements vs. pendulum based measurements. Schematic of projected force onto the measurement direction in a typical cantilever experiment (Backholm & Bäumchen, 2019; Kamimura & Takahashi, 1981; Kishino & Yanagida, 1988) versus a pendulum setup. Applied forces (**F**_*xy*_) shown as red arrows, measured force projections (**F**_*m*_) in gray. (**A**) Force projection schematic of a cantilever force measurement setup. Measured forces are 1D projections of applied forces on the measurement axis (**F**_*m*_ = **F**_⊥_). (**B**) Pendulum force measurement setup. Deflection direction is parallel to the horizontal projection of the applied force (**F**_*m*_ = **F**_*xy*_).

Here we propose a method that enables measurements of 2D horizontal component (Fig. 1B) of sub-mN forces applied by an untethered moving plant stem. Our method is based on the deflection of a free physical pendulum by the stem movement. This simple setup is cost-effective and modular, and can be adjusted to fit multiple scales of force and sample dimensions. As a case study, we measured forces applied by circumnutating bean plants prior to twining.

## Materials and Methods

### Plant materials and growth conditions

We measured forces applied by the climbing bean plant (*Phaseolus vulgaris*) prior to twining. Seeds of the Helda strain were purchased from Yarok organic nursery (https://www.ecolution.co.il/). Soil was acquired from Manna Center’s Program for Food Safety and Security at Tel Aviv University and mixed with sand (30%). Seeds were placed on a 1 cm thick piece of cotton wool spread out in a lidded flat box, kept within a growth chamber (Percival; Plant Tissue Culture; CU-41L5), at 24 °C. Seeds were kept moist and in darkness until germinated roots were roughly 1 *−* 3 cm long. Seedlings were removed from the growth chamber and planted in 100 ml soil pots with 25 *±* 5 Wm^*−*2^ instensity light for 16 hours a day. When stems reached 7 *−* 10 cm, plants were transferred to 1 L pots under 8 *±* 1 W m^*−*2^ intensity lighting, and watered once with 2 ml of 20-20-20 N-P-K fertilizer of 2% on the day of transfer to the larger pot. Thereafter watering was of 40 ml every two days, and the temperature kept at 24 °C. Experiments were done under constant LED lighting of 8 *±* 2 W m^*−*2^. All lamps were LED IP65 40 W, 4000 K, and light intensities were measured over a waveband between 300 *−* 1000 nm.

### Pendulum force measurement setup

The method developed in this paper is based on the deflection of a pendu-lum. The pendulum is positioned at such a distance from the plant, that the stem will encounter the rod during its circumnutation motion (Fig. 2). The position of the rod, and the contact position of the stem along it, are tracked by two cameras, positioned orthogonally on tripods, one from the side and one above the rod (see Fig. 2 and Fig. 3). The rod deflection is tracked over time and translated to force trajectories as elaborated in subsequent sections. The specifications of the setup components can be adjusted according to specific experiment requirements. In our setup, the pendulum rod is suspended from a structure (Ohad & Meroz, 2025) (hereafter: “add-on structure”) attached to the top-view camera lens. This structure (see Fig. 2) comprises of a circular piece, connected with M3 screws to two posts, which extend down vertically from opposite sides. A string (0.25 mm thick) connects the *≈* 9 cm wide gap between the posts at their lower end. The string is tightened and wrapped around the posts for stability.

**Fig. 2.**
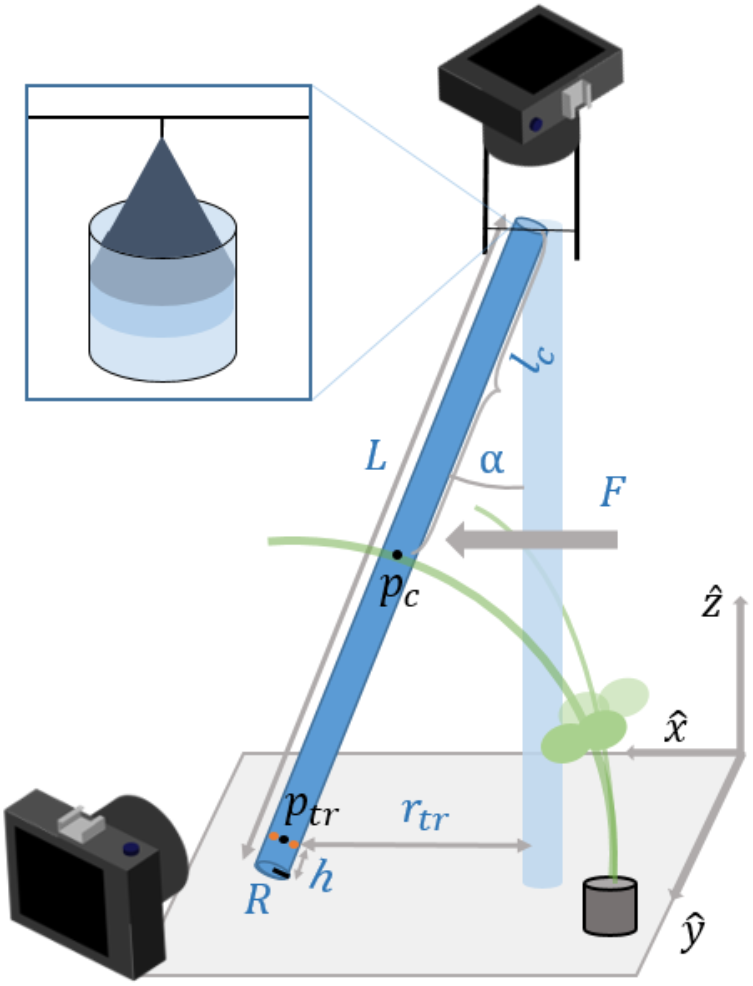
Schematic of experimental measurement setup. Cameras are placed at top and side positions with respect to the pendulum. Lens add-on structure - extending two posts with a string between them, is attached to the top-view camera, to which the pendulum rod is attached via the cone adapter (see inset). The vertical light blue bar represents the pendulum rod at rest before contact, and the solid blue bar depicts the rod position after a force is applied by the stem. A plant is positioned near the rod (light green: contact with the rod begins; solid green: some time after contact begins). Parameters include full pendulum length (*L* - includes cone adapter) and radius (*R*), distance of contact point from hinge (*l*_c_), deflection angle (*α*), distance of the tracked point (*p*_tr_) from the rod tip (*h*), distance (*d*) of the stem tip from contact point (*p*_c_), the horizontal distance of the tracked point from the vertical axis (*r*_tr_) and the applied force (*F*). The inset shows a close-up of the connection between the rod and the string mediated by the cone adapter; the centralizing 3D-printed cone is glued within the top of the rod. 3D axis directions shown on bottom right.

**Fig. 3.**
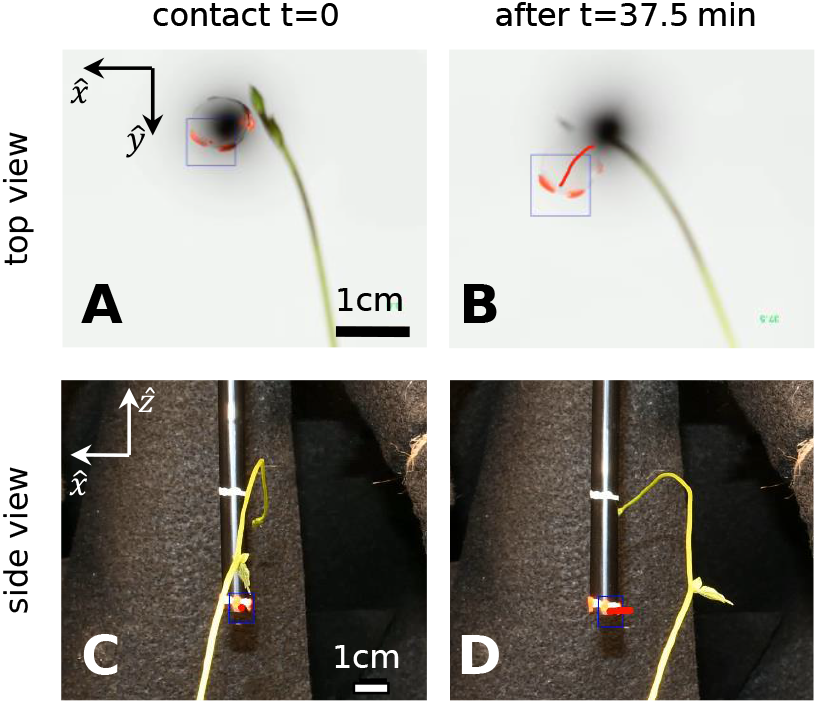
Top and side view tracking. (**A**,**B**) Top view, tracking box (blue) around the dye drop (orange) on the edge of the cylinder (thin black circle). (**A**) Contact initiation with the stem. The rod shadow is within the rod perimeter, indicating verticality. The stem impacts the rod (green) and in (**B**) is the rod position after 37.5 minutes, with the tracking history drawn (red points). Size and axis references are on the bottom right and top left respectively. (**C**,**D**) Side view depiction for the same event as in (A,B). Rod (black rod with white stripe) with tracking markers on the bottom (orange) and the tracking box in blue. Size and axis references are on the bottom right.and top left respectively

The plant base was positioned at a distance that ensures contact by placing the rod within reach of circumnutation cycles. At the start of each experiment, a 30 s interval time lapse is initiated simultaneously on both cameras.

Timelapse for both views was started simultaneously, manually, at the start of the experiment, incurring a maximal delay of half a second between views, negligible compared to the plan movement timescale. If the setup is used to measure faster dynamics, a wired intervalometer should be used to ensure higher synchronization between views.

If the stem slipped from the rod, the next circumnutation would typically lead to contact at a different position, which is treated as a separate event (the rod reaches equilibrium after approximately a quarter of the natural period, which is 1*/*4 s for the rod lengths we use – see detailed calculation in the SM.)

#### Pendulum setup

To enable the rod to be suspended from the camera add-on structure, we glued a 3D-printed adapter cone (Ohad & Meroz, 2025) within the top of the hollow rod (see inset in Fig. 2) and inserted a string through a hole in the top of the cone, attaching it to the horizontal string above it. The rod tip was typically placed about 1 cm below the stem tip height to ensure contact was made during the motion. To ensure the rod hangs straight, a 3D printed cone adapter was glued within the upper part of the rod (a plastic drinking straw). A string is pulled through a hole in the cone, allowing us to hang the cone (and the rod with it) on the horizontal string.

We designed a different cone for non-hollow rods (wood rods), We 3D-printed a small hollow cone to fit onto the wooden rod (as opposed to being placed inside the hollow plastic rods), see Figure S4, with further details in the SM. We level the cameras using bullseye levels on the tripods, and confirm the rod is vertical in both the top view (where the rod should appear as a dark circle - see Fig. 3A), and the side view (Fig. 3C).

Since the plant is not tethered to the device, the stem may contact the rod in suboptimal positions, i.e. near the bottom leading to slipping, or the top end, close to the hinge (where deflections are small with dominant errors). To avoid this, we adjusted plant and camera positions in advance. Furthermore, while slippage may be mitigating by increasing the friction of the rod, some slipping is required in order to allow the shoot apex to slip past the rod and thus twine.

#### Camera setup

Both top and side-view cameras are Nikon D7500 models, with a resolution of 5568×3712 pixels, pow-ered by a Glorich EP-5B DC coupler EN-EL15 EN-EL15a Dummy Battery EH-5 AC Power Supply Adapter Power Connector for Nikon. The top-view camera is mounted on a Manfrotto mk055 xpro3 tripod face down, with a Tamron lens 24 *−* 70 *f/*2.8 Di g2 macro lens attached. The side-view camera uses a Nikon AF-S DX NIKKOR 18 *−* 140 mm *f/*3.5 *−* 5.6G ED VR, supporting a wider field of view, and is fixed on a Joby tripod: GorillaPod 3K Kit for orientation and stability. The focus of the top view camera is fixed on the tip of the pendulum rod. The distance of the top lens from the lower tip of the rod was around 30 cm allowing supports of different lengths. The side view camera was positioned with the lens 80 cm away from the rod. Top (side) photos are taken against a white (black) background to increase contrast.

### Translating displacements into force trajectories

#### Tracking rod displacement over time

We place a drop of fluorescent dye (Ferrario, colore acrilico, rose pink) near the bottom end of the rod visible from both camera views (Fig. 3). We track the position of the dye (define as point *p*_tr_), and the contact position of the stem with the rod (*p*_c_) from the side view (see schematic in Fig. 2).

We note that the tracked point was chosen arbitrarily, since the value of interest is the difference between the starting and current point, rendering the particular tracked point irrelevant for the measurement.

Tracking uses CV2 (Bradski, 2000) library, utilizing a CSRT tracking algorithm. We use a script allowing selection of a region of interest, and tracks it over successive images in a folder. Returns coordinates of the upper left corner with the height and width of the tracking box per image (algorithm changes box properties per frame). Manual corrections are possible and were used as needed when occlusions or rapid motions caused the misplacement of the track box relative to the target object. Manual corrections to the tracked contact position *p*_c_ are applied when the tracking algorithm is displaced due to occlusions, rapid motion, or contrast issues. To convert distances measured in pixels into physical lengths, we use the known rod dimensions for reference, as well as a black and white checker pattern.

From the 2D projections of the tracked points (*p*_tr_ and *p*_c_) from the top and side views, we derive their respective 3D coordinates (Fig. 2). Based on these values, basic trigonometric arguments allow the extraction of the deflection angle of the pendulum *α* (see Fig. 2) by following (see SM for details):

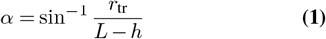

where *α* is the angle between the rod and the vertical, *r*_tr_ is the horizontal displacement of the tracked point *p*_tr_ from the starting position, *L* is the total length of the pendulum and *h* is the distance between *p*_tr_ and the rod tip, as defined in Fig. 2). Fig. 4A shows an example of the resulting angle trajectory *α*(*t*), based on measured deflections.

**Fig. 4.**
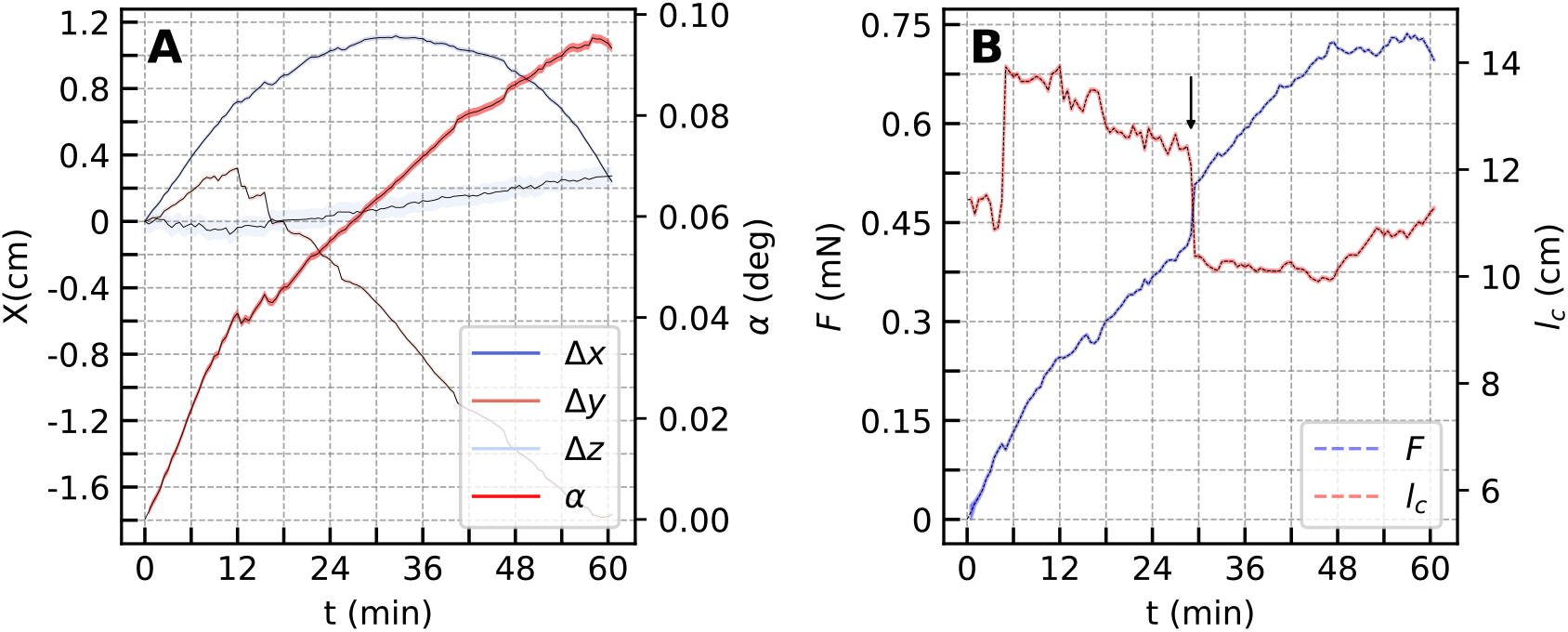
Circumnutation driven deflections, extracted angle, contact position and calculated force with error estimations. (**A**) Tracked coordinates Δ*x* = *x*(*t*) − *x*(*t* = 0) (and likewise for *y, z* coordinates) result from the generated force. The angle *α*, calculated by the rod tip deflection relative to the rest position, for one example event (red). (**B**) Distance of the contact point from pendulum hinge, *l*_c_ (dashed red), and the resulting calculated force *F* (dashed blue), for the trajectory in (A). Sharp change of *l*_c_ due to jumps of the shoot along the rod may result in shifts in the calculated force. An example at 30 min (black arrow). Colored shaded areas represent error estimations for coordinates, angle, *l*_tip_ and the extracted force (see Equations 3, 4, S.16 and S.18).

#### Torque equilibrium

The force applied by the plant stem is calculated via a torque equilibrium equation (Morin, 2008) of the form:

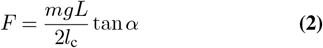

where *m* is the pendulum mass, *g* is the gravitational acceleration, *l*_c_ is the distance of the stem-contact point *p*_c_ (see Fig. 2) to the hinge point, and the angle *α* is calculated from Eq. 1. A detailed derivation can be found in the SM.

### Pendulum specifications

The basic component used for our setup is a hollow plastic tube (a plastic drinking straw). As detailed in the Results section, changing the rod dimensions and mass alters the range of force which can be measured, as well as the system sensitivity.

We note that the radius of the rod increases the pendulum-plant contact area. Furthermore, a circular cross-section, characterized by radial symmetry, is required in order to avoid torques around rotating the rod around its long axis, with adverse effects on the force measurements.

We produced pendulum rods of different masses and lengths for use in our experiments (see Table 2). To change the mass of the rod without changing the length, we 3D-printed hollow cylinders (Ohad & Meroz, 2025) with an outer diameter and length slightly smaller than those of the basic rod; we then inserted one cylinder into the basic rod. Changing the density and inner diameter of the printed cylinders give further flexibility in setting the rod mass while retaining rod length. Cylinders (and other 3D printed objects) were printed in a Sindoh 3DWOX DP200 3D printer, with ELEGOO PLA Filament, 1.75 mm diameter. For lighter rods we first use a slightly shortened basic rod and also produced a lower-mass rod using a Balsa wood beam smoothed down with sandpaper (see Table 2).

**Table 2.** Table of used rod parameters. Mass (*m*) includes cone adapter mass. *L* is the total pendulum length, *R* the rod radius, *α*_min_ is the minimal detectable angle, *F*_min_ the minimal detectable force and *F*_max_ the maximal force, calculated assuming a maximal deflection of 15°, with a contact position at a quarter of the pendulum length. The effective force range can be extended in either direction, see main text.

| $m$ (g) | $L$ (cm) | $R$ (cm) | $\alpha_{\min}$ ($^{\circ}$) | $F_{\min}$ ($\mu N$) | $F_{\max}$ (mN) |
| --- | --- | --- | --- | --- | --- |
| 0.20 | 19.9 | 0.2 | 0.04 | 0.5 | 1.1 |
| 0.65 | 17.7 | 0.8 | 0.04 | 2.0 | 3.4 |
| 0.86 | 19.9 | 0.8 | 0.04 | 2.3 | 4.5 |
| 1.10 | 19.9 | 0.8 | 0.04 | 3.0 | 5.8 |
| 3.10 | 19.9 | 0.8 | 0.04 | 8.4 | 16 |

## Results

### Calibration

To calibrate the system, we apply predetermined forces to the rod by using test weights (Acuris instruments) within the range of expected forces for the plant (from trial experiments). Weights were attached to a point on the pendulum, and compared to the extracted measured force. Using a pulley, the weights apply a known horizontal force equal to their weight (Fig. 5A inset). Fig. 5A presents the calculated force using Eq. 2 from the pendulum deflection, plotted against the known test weights, expecting a 1:1 relation for a calibrated system. The torque equilibrium calculation takes the horizontal component of the applied force into account.

**Fig. 5.**
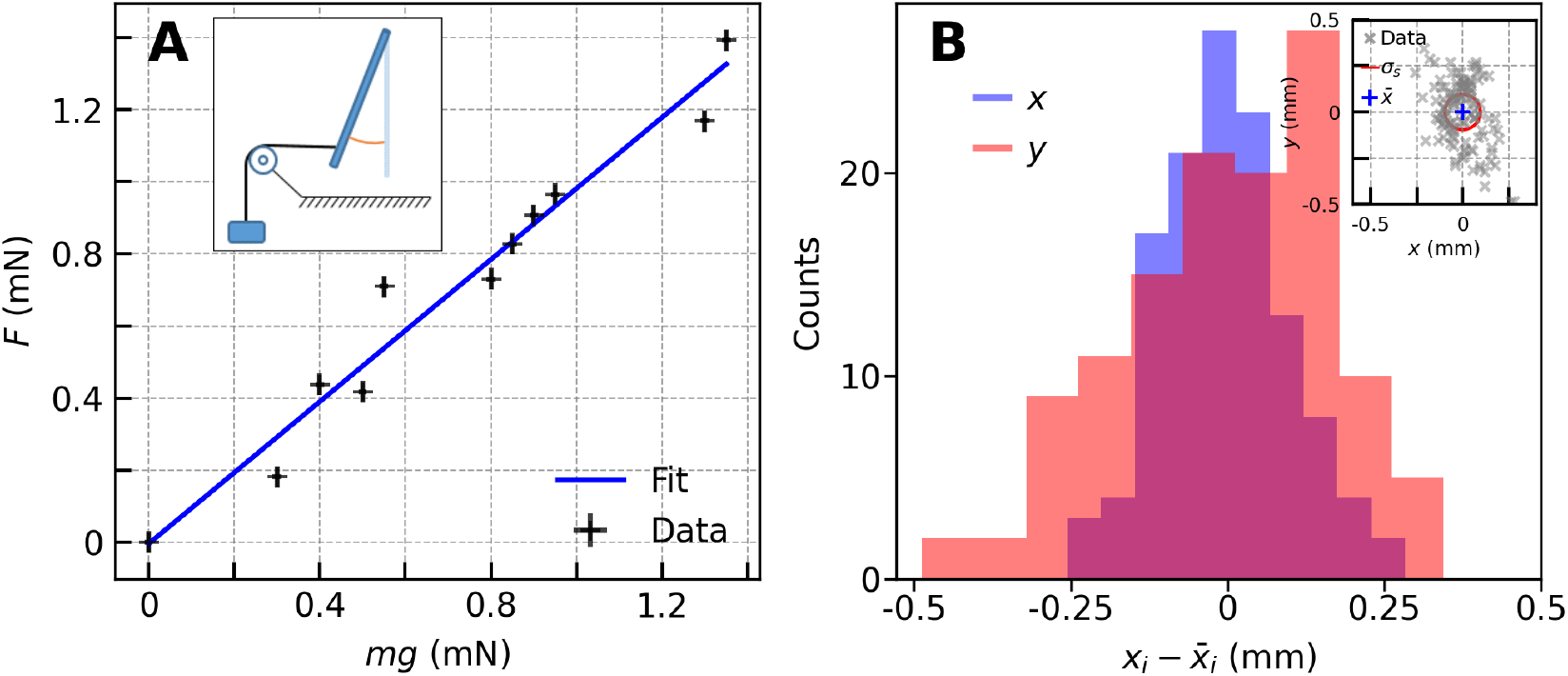
Calibration and noise. (**A**) Comparison of the calculated measured force (*F*) using our setup, for scaled measured weights (*mg*). Inset: Calibration setup schematic. A force is applied to the rod by a known mass, at a known position, to the rod via a pulley. We expect a 1:1 ratio between the values, as the applied force is designed to be applied horizontally. Goodness of fit parameters are *R*^2^ = 0.96. (**B**) Distribution of x_*i*_ coordinates (x and y) distance from mean position 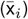 for top view tracking of inherent noise associated with the pendulum position with no directed forces applied. Inset: Tracked rod positions (gray x markers) and standard deviation of absolute distance (*σ*_n_ *≈* 0.1 mm red circle) relative to the mean position 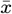 centered at *x* = 0 and *y* = 0 (blue +).

We note that rods with extreme slenderness ratios or stiffnesses may exhibit deformations. The maximal deformation is bounded by the deformation due to self weight of a horizontally place rod. Notably, the calibration process takes care of this.

### Force trajectories

Fig. 4 displays an example of the measurement of a force trajectory, representing the force a bean plant applied on the pendulum rod during its inherent circumnutation movement. Fig. 4A shows the tracked 3D coordinates of the contact point between the bean shoot and the pendulum, as well as the calculated deflection angle of the rod *α* with their respective errors. See Fig. S5 in SM for an overlaid image of the stem and support at different time points.

Fig. 4B shows the resulting force trajectory, following the computation described in the Methods. We ran experiments on *N* = 50 plants (10 per rod mass), measuring the forces they applied on the pendulum setup during their circumnutation movement. Fig. 6A shows the resulting force trajectories. The trajectories exhibit a characteristic increase in force until either the stem slips from the pendulum, where we allow a new measurement after a circumnutation period, or it starts to twine on the rod - at both points the measurement is stopped.

**Fig. 6.**
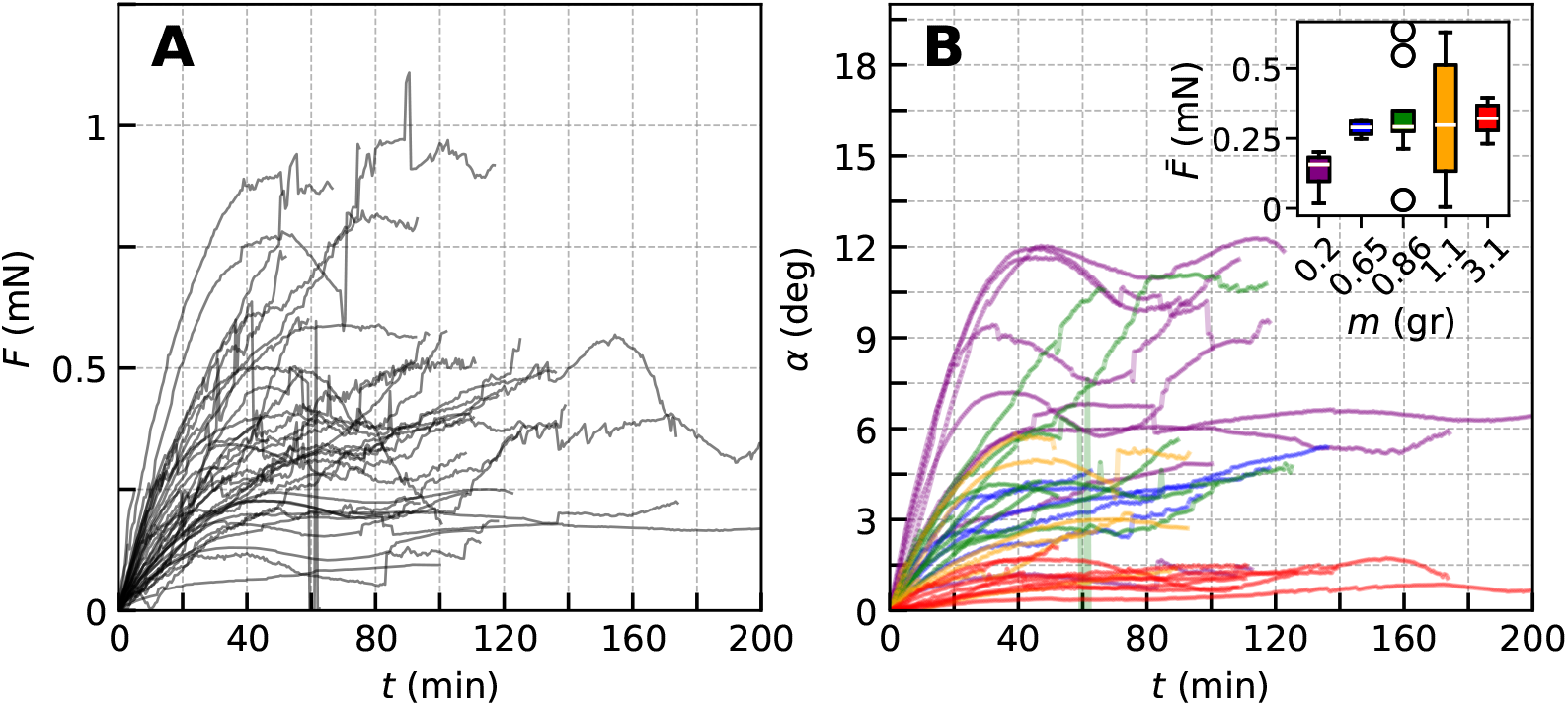
Force trajectories and corresponding deflection angles per rod mass for 2-3 week old *Phaseolus vulgaris* stems, *N* = 50. (**A**) All force trajectories (*F*) measured with the force measurement setup after calibration. (**B**) Deflection angles (*α*) for rods of different mass (by color). The maximal deflection depends noticeably on the rod mass, allowing for varied force resolution and measurement range. In the inset, distributions of the mean force 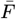 per support mass, with the line in the box as the median, the upper and lower box edges as the first and third quartiles, and the whiskers showing the minimal and maximal values within 1.5 times the inter-quartile range of the box edges.

### Rapid force variations

We may observe two types of abrupt changes in the measured force trajectories. The first is due to sliding of the stem along the rod, represented by jumps in the tracked contact position *p*_c_. Such motions may be large (several cm) and fast (seconds), and may result in rapid shifts in the force (see example in Fig. 4B). Additionally, the initial contact with the stem may cause the rod to bounce, thus briefly losing contact. These gaps are much faster than the temporal resolution in our experiments, and therefore negligible.

### Errors and Resolution

The temporal resolution is dictated by two parameters. First, the time between pictures in the timelapse photography, here set at 30 seconds, was chosen to allow a smooth coverage of the shortest recorded trajectories (shortest time until contact termination). The second parameter is the relaxation time of the rod when it is bumped during the first encounter with a circumnutating plants. This can be estimated as roughly a quarter of the natural period of the pendulum, equal to *∼* 1*/*4 sec (calculation details in SM). Therefore, significantly faster movements can be measured with this setup.

The image resolution set by the cameras is 1 pixel, equivalent to 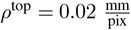 and 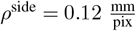 in the top view and side view respectively. The tracking algorithm introduces some uncertainty, which is estimated as *σ*_tr_ *≈* 5 pixels, which results in an effective resolution of *≈* 0.1 mm in the top view.

The inherent noise of the system is defined as the standard deviation of fluctuations in the position of a free rod measured over 2 minutes, at 1 s intervals (Fig. 5B). The fluctuations from this noise amount to *σ*_n_ = 0.1 mm, which is comparable to the effective tracking resolution. We define the total error of any tracked point *σ*, taking into account the said contributions, following:

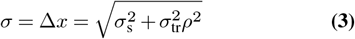

where *σ*_*tr*_ is the tracking error in pixels, *σ*_*n*_ is the standard deviation of the inherent fluctuations and *ρ* is the mm value of one pixel (different for top and side views). Based on these errors we can estimate the error in the calculated deflection angle according to (see SM for derivation):

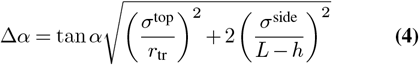

where *σ*^top^ and *σ*^side^ are the total coordinate errors for the top and side views respectively following Eq. 3 with *ρ* changing according to view. The error in the force measurements is mostly affected by the position error, see the full expression for the force error in the SM (Eq. S.18).

### Tuning the range of measurable forces

The sensitivity of the force measurement setup, i.e. the minimal force that can be detected, is dictated by the minimal deflection angle that can be detected in the system, namely:

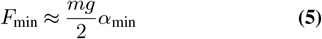

where *m* is the mass of the rod, *g* is the gravitational acceleration. In turn, the angle sensitivity *α*_min_ is dictated by the position resolutions *σ* (see Eq. 3), following Eq. S.21, obtained by assuming the minimal detectable deflection *σ*^top^ and contact near the rod tip (see SM for derivation).

Therefore, for a given image resolution *ρ*, the sensitivity of the setup can be tuned by choosing different configurations of the pendulum rod, namely rod mass *m* and length *L*, thus allowing measurements of a wide range of forces. This is demonstrated in Fig. 6B where, for similar forces, smaller rod masses result in larger deflection angles *α*. As angle resolution *α*_min_ is system dependent, weaker forces will require lighter rods, ensuring the deflection remains significant compared to the error.

The maximal measurable force is given by the maximal deflection angle, as dictated by a number of parameters such as the camera field of view, or shoot slippage. In our setup we observed a maximal angular deflection of *α*_max_ *≈* 12°. Therefore if forces are such that deflection angles are too large, a heavier rod is required.

For example, a 50 gr, 100 cm rod would have a force detec-tion range of approximately 0.03 *−* 260 mN, while a 0.1 gr, 10 cm rod could measure forces in a range of 0.3 *−* 300 *µ*N. Force ranges are provided in Table 2 for a number of different rod configurations used in our measurements.

## Discussion

We have presented an optically based force measurement system utilizing a freely jointed physical pendulum allowing measurement along 2 axes (as opposed to commonly available 1 axis). This setup enables measurements of *µ*N forces applied by freely moving untethered organs or organisms without the use of force measurement load cells. It is particularly relevant for the measurement of growth-driven forces applied by growing plants on their environment. We described the physical setup, force calculation method, calibration and error limitations. Finally, we demonstrated how setup sensitivity and range can be modified by varying pendulum mass and length. This system addresses a need for a flexible tool to measure a wide range of forces exerted by plants of diverse dimensions. Indeed, most existing techniques for measuring *µ*N-mN scale forces are designed for organisms below the mm range, whereas our setup can be adapted to suit organisms on the meter scale. Crucially, unlike existing systems for measuring forces generated by plants, our system does not require tethering of the organism to the measurement device while also producing 2D force measurements, enabling new directions for testing plant behaviors and interactions.

### Limitations

Our setup sensitivity is limited by the optical resolution of the cameras, and the inherent noise of the system. The temporal resolution is set by the timelapse interval, as well as by camera capabilities. Since our force extraction method relies on the applied force being primarily horizontal, forces with a significant vertical component will be underestimated using this method.

The absence of tethering may also pose a challenge as fluctuations in the motion of the organism may not guarantee contact with the rod, requiring specific adaptations per test case. Slippage may be decreased by increasing the friction coefficient between the rod and sample, e.g. by using a rough rod, however this will limit the inherent movements of the sample, and may limit the movement of the apex past the rod, required for twining.

If no friction is added, the stem may contact the rod in sub-optimal positions, i.e. near the bottom leading to slipping, or the top end, close to the hinge (where deflections are small with dominant errors). As such, plant and camera placements and heights should be adjusted to avoid sub-optimal contact position.

### Possible applications

Our system allows non-invasive investigation of mechanical interactions between organisms and their environment. It may facilitate the estimation of forces applied by leaf climbers prior to substrate adhesion (Bauer et al., 2010; Steinbrecher et al., 2010), as well as forces involved in posture control by stem twiners (Onyenedum et al., 2025) during growth-driven movements, etc. With appropriate scaling, larger setups could be adapted to measure forces in liana stems, such as those of grapevines and wisteria (Onyenedum & Pace, 2021; Hattermann et al., 2022). Additionally, low mass supports may allow the detection of pushing and pulling forces exerted by small non-plant organisms which may be difficult to measure with traditional setups - for example, by coating the rod tip with sugar and measuring forces generated by different insects (Feinerman et al., 2018).

## Conclusion

This setup provides 2D force measurements without requiring physical attachment of the organ to the device. This system provides unique opportunities for exploring mechanical interactions between plants and their environments, especially when studying weak forces exerted by larger-scale biological samples, such as growing stems and roots, which are typically not suitable for existing micro-scale measurement methods. Future refinement of the approach could further extend its applicability to explore interactions across diverse biological species and length scales.

## Acknowledgements

Y.M. acknowledges support from the Israel Science Foundation Research Grant (ISF) no. 2307/22, and ERC grant GROWsmart 101165101. A.O. acknowledges support from the Colton Foundation scholarship.

We thank Roni Kempinski for assistance in implementing image analysis tools. We thank Alessia Perilli, Amir Porat, Mathieu Riviere and Agueda de la Vega, for consulting, assisting and troubleshooting.

## Author contributions

AO and YM designed the research. AO collected data and AO analyzed the data with guidance from YM. AO and YM wrote the manuscript.

## Data availability

We have put the full workflow for 5 example trajectories on a Zenodo repository (https://doi.org/10.5281/zenodo.15545548; (Ohad & Meroz, 2025)), including code and raw data. Also included are calibration and noise measurements.

Other experimental data available upon request.

## Supplementary data

The following supplementary data are available online.

Video. S1. Top and side views with force trajectory.

Fig. S1. 2D setup definitions schematic.

Fig. S2. 2D torque schematic.

## References

Darwin. The movements and habits of climbing plants. Nature, 13(317):65–66, November 1875.

Gianoli. The behavioural ecology of climbing plants. AoB PLANTS, 7, January 2015.

Moulia et al. The shaping of plant axes and crowns through tropisms and elasticity: an example of morphogenetic plasticity beyond the shoot apical meristem. New Phytologist, 233(6):2354–2379, 2022.

Tedone et al. Optimal control of plant root tip dynamics in soil. Bioinspiration amp; Biomimetics, 15(5):056006, July 2020.

Goriely. The Mathematics and Mechanics of Biological Growth. Springer, 2017.

Goriely & Neukirch. Mechanics of climbing and attachment in twining plants. Physical Review Letters, 97(18), November 2006.

Chelakkot & Mahadevan. On the growth and form of shoots. Journal of The Royal Society Interface, 14(128):20170001, March 2017.

Okada & Shimura. Reversible root tip rotation in <i>arabidopsis</i> seedlings induced by obstacle-touching stimulus. Science, 250(4978):274–276, 1990.

Thompson & Holbrook. Root-gel interactions and the root waving behavior of arabidopsis. Plant Physiology, 135(3):1822–1837, July 2004.

Zhang et al. A mechano-sensing mechanism for waving in plant roots. Scientific Reports, 12(1), jun 2022.

Porat et al. On the mechanical origins of waving, coiling and skewing in arabidopsis thaliana roots. Proceedings of the National Academy of Sciences, 121(11), March 2024.

Telewski & Jaffe. Thigmomorphogenesis: Field and laboratory studies of abies fraseri in response to wind or mechanical perturbation. Physiologia Plantarum, 66(2):211–218, February 1986.

Jaffe. Thigmomorphogenesis: The response of plant growth and development to mechanical stimulation. Planta, 114(2):143–157, 1973.

Biddington. The effects of mechanically-induced stress in plants ? a review. Plant Growth Regulation, 4(2):103–123, 1986.

Jaffe et al. Thigmo responses in plants and fungi. American Journal of Botany, 89(3):375–382, March 2002.

Braam. In touch: plant responses to mechanical stimuli. New Phytologist, 165(2):373–389, November 2004.

Forterre et al. How the venus flytrap snaps. Nature, 433(7024):421–425, January 2005.

Takahashi & Jaffe. Thigmotropism and the modulation of tropistic curvature by mechanical perturbation in cucumber hypocotyls. Physiologia Plantarum, 80(4):561–567, December 1990.

Moulia et al. Mechanosensitive control of plant growth: bearing the load, sensing, transducing, and responding. Frontiers in Plant Science, 6, February 2015.

González-Teuber & Gianoli. Damage and shade enhance climbing and promote associational resistance in a climbing plant. Journal of Ecology, 0(0), November 2007.

Kolb et al. Physical root–soil interactions. Physical Biology, 14(6):065004, November 2017.

Kolb et al. Root–soil interaction. In Soft Matter in Plants: From Biophysics to Biomimetics. The Royal Society of Chemistry, 09 2022.

Quiros et al. Plant root growth against a mechanical obstacle: the early growth response of a maize root facing an axial resistance is consistent with the lockhart model. Journal of The Royal Society Interface, 19(193), August 2022.

Koren et al. Analysis of root-environment interactions reveals mechanical advantages of growth-driven penetration of roots. Plant, Cell amp; Environment, 47(12):5076–5088, August 2024.

von Sachs. Physiology of plants. Clarendon Press, Oxford, 1887.

Silk. On the curving and twining of stems. Environmental and Experimental Botany, 29(1):95–109, January 1989.

Stolarz. Circumnutation as a visible plant action and reaction. Plant Signaling & Behavior, 4(5): 380–387, May 2009.

Bai et al. Behavioral and economic traits reflect distinct resource acquisition strategies in tendril vines and stem twining vines. Ecology and Evolution, 14(9), September 2024.

Rivière et al. Plant nutation relies on steady propagation of spatially asymmetric growth pattern. Quantitative Plant Biology, pages 1–25, 2025.

Jaffe. Reversible force transduction in tendrils of passiflora coerulea. Plant and Cell Physiology, 11(1):47–53, February 1970a.

Silk & Hubbard. Axial forces and normal distributed loads in twining stems of morning glory. Journal of Biomechanics, 24(7):599–606, January 1991.

Matista & Silk. An electronic device for continuous, in vivo measurement of forces exerted by twining vines. American Journal of Botany, 84(8):1164–1168, August 1997.

Silk & Holbrook. The importance of frictional interactions in maintaining the stability of the twining habit. American Journal of Botany, 92(11):1820–1826, November 2005.

Steinbrecher et al. Quantifying the attachment strength of climbing plants: A new approach. Acta Biomaterialia, 6(4):1497–1504, April 2010.

Bauer et al. Always on the bright side: the climbing mechanism of galium aparine. Proceedings of the Royal Society B: Biological Sciences, 278(1715):2233–2239, December 2010.

Klimm et al. Force generation in the coiling tendrils of passiflora caerulea. Advanced Science, 10 (28), August 2023.

Klimm et al. Natural coil springs: Biomechanics and morphology of the coiled tendrils of the climbing passion flower passiflora discophora. Acta Biomaterialia, 189:478–490, November 2024.

Jaffe. Physiological studies on pea tendrils. Plant Physiology, 45(6):756–760, June 1970b.

Kamimura & Takahashi. Direct measurement of the force of microtubule sliding in flagella. Nature, 293(5833):566–568, October 1981.

Kishino & Yanagida. Force measurements by micromanipulation of a single actin filament by glass needles. Nature, 334(6177):74–76, July 1988.

Beaussart et al. Quantifying the forces guiding microbial cell adhesion using single-cell force spectroscopy. Nature Protocols, 9(5):1049–1055, April 2014.

Backholm & Bäumchen. Micropipette force sensors for in vivo force measurements on single cells and multicellular microorganisms. Nature Protocols, 14(2):594–615, January 2019.

Poppele & Hozalski. Micro-cantilever method for measuring the tensile strength of biofilms and microbial flocs. Journal of Microbiological Methods, 55(3):607–615, December 2003.

Autumn et al. Adhesive force of a single gecko foot-hair. Nature, 405(6787):681–685, June 2000.

Doll et al. Su-8 force sensing pillar arrays for biological measurements. Lab on a Chip, 9(10): 1449, 2009.

Sznitman et al. Propulsive force measurements and flow behavior of undulatory swimmers at low reynolds number. Physics of Fluids, 22(12), December 2010.

Rabets et al. Direct measurements of drag forces in c. elegans crawling locomotion. Biophysical Journal, 107(8):1980–1987, October 2014.

Das et al. The biomechanics of the locust ovipositor valves: a unique digging apparatus. Journal of The Royal Society Interface, 19(188), March 2022.

Filc et al. Tailoring the mechanical properties of high-fidelity, beetle-inspired, 3d-printed wings improves their aerodynamic performance. Advanced Engineering Materials, 25(21), September 2023.

Ohad & Meroz. Data and code for: Dual-axial force measurements of non-tethered plants. 10.5281/zenodo.15545548, 2025. Developed by Amir Ohad.

Bradski. The OpenCV Library. Dr. Dobb’s Journal of Software Tools, 2000.

Morin. Introduction to Classical Mechanics: With Problems and Solutions. Cambridge University Press, 2008.

Onyenedum et al. Gelatinous fibers develop asymmetrically to support bends and coils in common bean vines (phaseolus vulgaris). American Journal of Botany, 112(3), March 2025.

Onyenedum & Pace. The role of ontogeny in wood diversity and evolution. American Journal of Botany, 108(12):2331–2355, December 2021.

Hattermann et al. Mind the gap: Reach and mechanical diversity of searcher shoots in climbing plants. Frontiers in Forests and Global Change, 5, April 2022.

Feinerman et al. The physics of cooperative transport in groups of ants. Nature Physics, 14(7): 683–693, May 2018.

Ku. Notes on the use of propagation of error formulas. Journal of Research of the National Bureau of Standards, Section C: Engineering and Instrumentation, 70C(4):263, October 1966.

Landau. Mechanics, Third Edition: Volume 1 (Course of Theoretical Physics). Butterworth-Heinemann, 3 edition, January 1976.

