## Supplementary for "Camera-Based Bi-Axial Measurement of Weak Forces Generated by Freely-Moving Plant Organs"

### Supporting information

Supplementary materials include one PDF file and one video as well as example data for 5 events.

**SM.pdf**: Contains an appendix detailing the derivation of reported values and associated error.

**Supplementary video**: Demonstrated tracking and development of the force trajectory over time.

**Data and code availability**: Datasets and analysis scripts used to generate trajectories are openly available via Zenodo: `https://doi.org/10.5281/zenodo.15545548`.

The Zenodo repository includes:

**Python scripts**: Auxillary python file and Jupyter notebook file for importing data and calculating deflections, angles and force trajectories pf example events.

**Excel spreadsheet**: Contains event data for use in analysis script.

**Tracked coordinates**: Top and side-view tracking data for the tracked point ($p_{\mathrm{tr}}$), both for contact events and non-contact fluctuations, as well as tracking of the stem's contact point with the rod ($p_{\mathrm{c}}$).

**Example datasets**: Images of top and side views for 5 example trajectories.

**3D-printable STL files**: Design files for lens add-on structures.

**README file**: Detailing folder contents.

**3D coordinate extraction.** We define three relevant points along the pendulum rod as vectors relative to a reference point (Fig. S2): the point of contact between the shoot and the rod $p_{\mathrm{c}}$ (whose coordinates can only be tracked from the side view - yielding coordinates $x_{\mathrm{c}}$ and $z_{\mathrm{c}}$), the fluorescent marker near the rod tip $p_{\mathrm{tr}}$ (which can be tracked from both top and side views yielding $p_{\mathrm{tr}} = (x_{\mathrm{tr}}, y_{\mathrm{tr}}, z_{\mathrm{tr}})$), the tip of the rod $p_{\mathrm{tip}}$. We obtain $p_{\mathrm{tip}}$ from the side view in the first frame of the time lapse, but do not track it over time. We define the constant distance between the tracked marker and the rod tip:

$$h \equiv |p_{\mathrm{tr}} - p_{\mathrm{tip}}| \tag{S.1}$$

The hinge point ($p_{\mathrm{h}}$), which is obtained by adding the pendulum length (excluding $h$) to the z value of the tracked point at rest:

$$p_{\mathrm{h}} = p_{\mathrm{tr}}(t=0) + (0,0,L-h)$$

Where the points are in the form of coordinates $(x,y,z)$ and

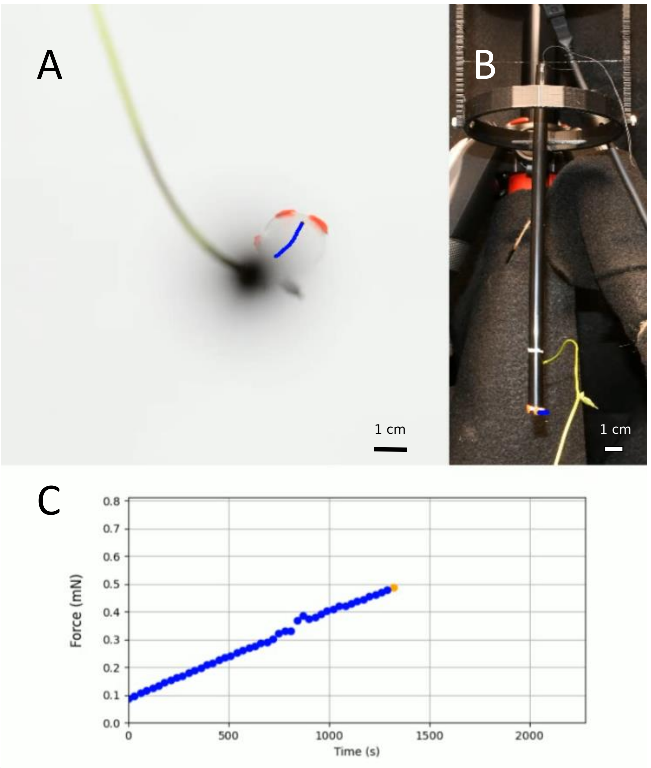


**Fig. S1. Top and side views with force trajectory.** Snapshot from video showing side (A) and top (B) views during contact with a 0.86 g rod. (C) Force trajectory developing over time. Scale bars at bottom of the images.

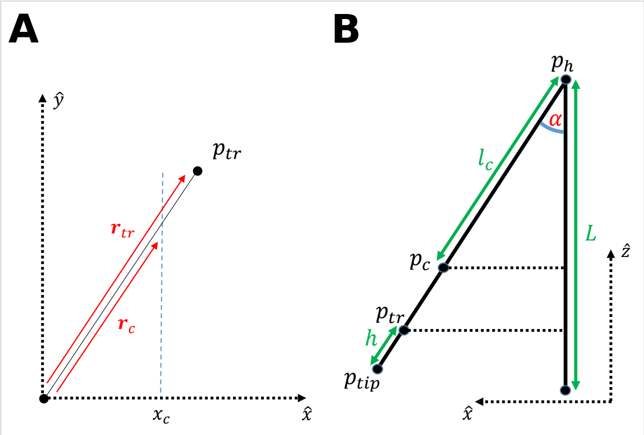


**Fig. S2. 2D setup definitions schematic**. (**A**) Top view: x,y projection of hinge (dot at origin), tracked ($p_{\mathrm{tr}}$) and contact ($p_{\mathrm{c}}$) points, with the corresponding vectors pointing to them: $\mathbf{r}_{\mathrm{tr}}, \mathbf{r}_{\mathrm{c}}$ (in red). Side view: (**B**) x,z projection for lengths and points in the setup. The tracked point ($p_{\mathrm{tr}}$) on the support is separate from the contact point ($p_{\mathrm{c}}$) and the rod tip ($p_{\mathrm{tip}}$). When the pendulum is at the rest point, we identify the distance of the track point from the tip of the rod ($h$). $L$ is the total length of the pendulum, and $lc$ is the distance between the pendulum hinge ($p_{\mathrm{h}}$) and the contact point ($p_{\mathrm{c}}$).

$L$ is the total length of the pendulum (Fig. S2). As the rod is deflected by the stem, we determine the deflections in the x,y plane with the top view camera:

$$r_{\mathrm{tr}}^2 = \Delta x_{\mathrm{tr}}^2 + \Delta y_{\mathrm{tr}}^2 \quad \textbf{(S.2)}$$

where $r_{\mathrm{tr}} \equiv |\mathbf{r}_{\mathrm{tr}}|$ is the length of the horizontal vector $\mathbf{r}_{\mathrm{tr}}$, $\Delta x_{\mathrm{tr}}, \Delta y_{\mathrm{tr}}$ are the coordinate differences between the tracked position of the rod tip at a given time t, and the initial position $x_0, y_0$ of the tracked point $p_{\mathrm{tr}}$:

$$\Delta x_{\mathrm{tr}}(t) = x_{\mathrm{tr}}(t) - x_{\mathrm{tr}}(t=0) \quad \textbf{(S.3)}$$

and likewise for the y coordinate of the tracked point. Next we extract the deflection angle by the trigonometric relation (1):

$$\alpha = \sin^{-1} \frac{r_{\mathrm{tr}}}{L-h} \quad \textbf{(S.4)}$$

where we omit the explicit time dependence of $\alpha, r_{\mathrm{tr}}$ for clarity.

Since the contact point can only be tracked from the side view (we cannot discern contact position reliably from the top view), we do not have the $y$ coordinate of the contact point $y_{\mathrm{c}}$. Instead we parameterize the support, using the hinge and tracked position to define a 3D vector. First we define the vector from the hinge to the tracked point:

$$\mathbf{V}_{\mathrm{tr}} = p_{\mathrm{h}} - p_{\mathrm{tr}}$$

To obtain the 3D position of the contact point $p_{\mathrm{c}}$, we first define the vector of the contact point:

$$\mathbf{V}_{\mathrm{c}} = p_{\mathrm{h}} - p_{\mathrm{c}}$$

Now, due to $\mathbf{V}_{\mathrm{c}}$ being a segment of $\mathbf{V}_{\mathrm{tr}}$, we use the linear parametric relations along $\mathbf{V}_{\mathrm{tr}}$ to obtain the contact position coordinates of $p_{\mathrm{c}}$:

$$k \equiv z_{\mathrm{c}}/z_{\mathrm{tr}} \rightarrow \begin{cases} y_{\mathrm{c}} = k * y_{\mathrm{tr}} \\ x_{\mathrm{c}} = k * x_{\mathrm{tr}} \end{cases} \quad \textbf{(S.5)}$$

Where $(x,y,z)_{\mathrm{c}}$ are the coordinates of the contact point $p_{\mathrm{c}}$ from the side-view camera, and $k$ is the proportion constant between the coordinates of $\mathbf{V}_{\mathrm{c}}$ and $\mathbf{V}_{\mathrm{tr}}$.

To extract the distance of the contact point from the hinge we take an Euclidean distance:

$$l_{\mathrm{c}} \equiv |\mathbf{V}_{\mathrm{c}}| = \sqrt{x_{\mathrm{c}}^2 + y_{\mathrm{c}}^2 + y_{\mathrm{c}}^2} \quad \textbf{(S.6)}$$

Alternatively, from the length $|\mathbf{V}_{\mathrm{tr}}|$, and the proportion $k$, we can extract the length $l_{\mathrm{c}}$ from it:

$$k \equiv z_{\mathrm{c}}/z_{\mathrm{tr}} = \frac{|\mathbf{V}_{\mathrm{c}}|}{|\mathbf{V}_{\mathrm{tr}}|} \quad \textbf{(S.7)}$$

$$l_{\mathrm{c}} \equiv |\mathbf{V}_{\mathrm{c}}| = k|\mathbf{V}_{\mathrm{tr}}| \quad \textbf{(S.8)}$$

**Quasi-static approximation.** We use the assumption that the state of the system is approximately in equilibrium at every moment, meaning a quasi-statically applied force; the slow circumnutation rate ($T_{CN} \approx 100\frac{\mathrm{min}}{\mathrm{s}}$ for a typical plant in our experiment), is around three orders of magnitude slower than the pendulum period (a period of $T_{pend} \approx 1$ s), demonstrating the separation of timescales between the slow circumnutations and the fast mechanical relaxation, and we assume from now on the dynamics are quasi-static. Under this assumption, the torque applied by the plant to the rod is equal and opposite to the the torque due to the pendulum weight at every moment.

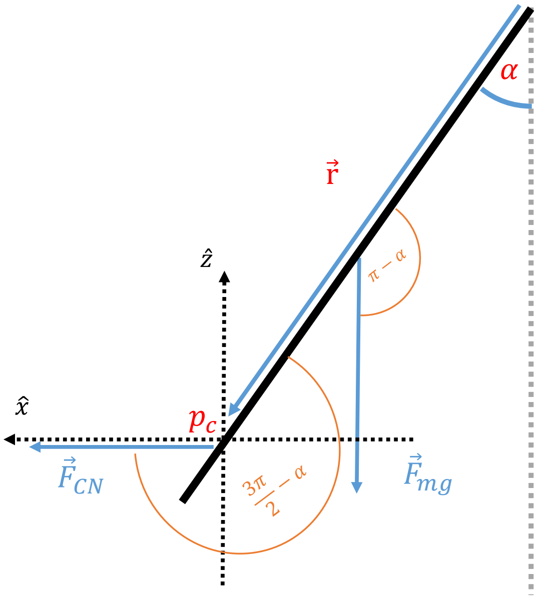


**Fig. S3. 2D torque schematic**. A force $F_{CN}$ is applied at the contact point $p_{\mathrm{c}}$, and the rod weight $mg$ applies a force in the negative z direction at half the rod length. Torque of the force is applied at $r$ from the origin. The torque generated by these forces reaches equilibrium when the pendulum is at an angle $\alpha$.

**Torque equilibrium.** To calculate the force $F$ required to deflect the pendulum to an angle $\alpha$ from the vertical, we calculate the torque applied by the self-weight of the pendulum, and assume this is equal to the torque applied by the force $F$ exerted by plant circumnutations at the point of contact $p_{\mathrm{c}}$ along the pendulum (see Fig. S3).

Torque, the rotational analogue of linear force, is given by

$\tau = \mathbf{r} \times \mathbf{F}$, where $\mathbf{F}$ is the force vector (applied by the plant), and $\mathbf{r}$ is the position vector (a vector from the point about which the torque is being measured - here the axis of rotation of the pendulum - to the point where the force is applied by the plant $p_\mathrm{c}$). For simplicity, we assume all forces and points are in the $\hat{\mathbf{x}} - \hat{\mathbf{z}}$ plane. In this case the magnitude of the torque associated with the force applied by the plant follows:

$$\tau_F = l_\mathrm{c} F \sin\left(\frac{3\pi}{2} - \alpha\right) \tag{S.9}$$

where $l_\mathrm{c}$ is the distance from the pendulum axis to the point of contact $p_\mathrm{c}$, and the sine function represents the cross product, with the angle between the pendulum vector $\vec{r}$ and the applied force vector $\vec{F}$ equal to $\sin\left(\frac{3\pi}{2} - \alpha\right) = -\cos(\alpha)$. Since we take the $\hat{x}$-axis in the opposite direction here, we can take the positive value of cosine. The magnitude of the torque applied by the self-weight of the pendulum follows:

$$\tau_W = \frac{L}{2} mg \sin(\pi - \alpha) \tag{S.10}$$

where the factor $\frac{1}{2}$ represents the center of mass in a uniform rod, and the sine function is again the angle between the pendulum vector $\vec{r}$ and the direction of gravity equal to $\sin(\pi - \alpha) = \sin(\alpha)$. Equating the torques in Eq. S.9 and Eq. S.10 we obtain the relation (Eq. 2):

$$F = \frac{mgL}{2l_\mathrm{c}} \tan\alpha \tag{S.11}$$

Substituting the deflection angle $\alpha$ (Eq. 1), and the distance to the contact point $l_\mathrm{c}$ from the hinge, yields the horizontal projection of the applied force $F$.

**Errors.** To compute the error per variable, we take the root of the sum of squares of partial derivatives for each parameter (Ku, 1966).

As mentioned in the calibration subsection, we take the resolution of tracking measurements $\sigma_{tr}$ to be 5 pixels in both side and top view, as there are small fluctuations during tracking of the marked points. The error for the coordinates values will be:

$$\sigma^\mathrm{top} = \Delta x^\mathrm{top} = \Delta y^\mathrm{top} = \sqrt{\sigma_\mathrm{n}^2 + \sigma_\mathrm{tr}^2 \rho_\mathrm{top}^2} \tag{S.12}$$

Where $\sigma_n$ is the standard deviation of the inherent fluctuations of the top view (see Fig. 5B) when no force is applied, and $\rho_\mathrm{top}$ is the cm value of 1 pixel in the top view.

Similarly, for the x,z axes from the side view we have:

$$\sigma^\mathrm{side} = \Delta x^\mathrm{side} = \Delta z^\mathrm{side} = \sqrt{\sigma_\mathrm{n}^2 + \sigma_\mathrm{tr}^2 \rho_\mathrm{side}^2} \tag{S.13}$$

Where $\sigma_{tr}$ and $\sigma_n$ remain the same, and $\rho_\mathrm{side}$ is the cm value of 1 pixel in the side view.

The error for the distance of a 3D vector from the origin (as in Eq. S.6), which was extracted from directly from the cameras, will depend on the view errors:

$$\Delta|\mathbf{V}| = \sqrt{\left(\frac{2x}{2|\mathbf{V}|}\Delta x\right)^2 + \left(\frac{2y}{2|\mathbf{V}|}\Delta y\right)^2 + \left(\frac{2z}{2|\mathbf{V}|}\Delta z\right)^2}$$
$$\Delta|\mathbf{V}| = \frac{1}{|\mathbf{V}|}\sqrt{(x\sigma^\mathrm{top})^2 + (y\sigma^\mathrm{top})^2 + (z\sigma^\mathrm{side})^2} \tag{S.14}$$

The error for the proportion constant $k$ from Eq. S.7 is given by:

$$\Delta k = \sqrt{\left(\frac{1}{z_\mathrm{tr}}\right)^2 \Delta z_\mathrm{c}^2 + \left(\frac{z_\mathrm{c}}{z_\mathrm{tr}}\right)^2 \Delta z_\mathrm{c}^2} =$$
$$\frac{\Delta z_\mathrm{c}}{z_\mathrm{tr}^2}\sqrt{1 + \left(\frac{z_\mathrm{c}}{z_\mathrm{tr}}\right)^2}$$
$$\Delta k = \frac{\sigma^\mathrm{side}}{z_\mathrm{tr}^2}\sqrt{1 + \left(\frac{z_\mathrm{c}}{z_\mathrm{tr}}\right)^2} \tag{S.15}$$

The error for the contact distance $l_\mathrm{c}$, from Eq. S.8:

$$\Delta l_\mathrm{c} = \sqrt{\Delta k^2 |\mathbf{V}_\mathrm{tr}|^2 + \Delta|\mathbf{V}_\mathrm{tr}|^2 k^2} \tag{S.16}$$

The compounded error for the angle $\alpha$, from Eq. 1, and using $\sin\alpha \equiv A = \frac{r_\mathrm{tr}}{L-h}$ we get:

$$\Delta\alpha = \sqrt{\frac{1}{\left(\sqrt{1-A^2}\right)^2}} *$$
$$\sqrt{\left(\frac{\Delta r_\mathrm{tr}}{L-h}\right)^2 + \left(\frac{\Delta L}{(L-h)^2}\right)^2 + \left(\frac{\Delta h}{(L-h)^2}\right)^2}$$

taking the square root from the left-most term and taking $A^2$ as a common factor from each fraction in the square root:

$$\Delta\alpha = \frac{A}{1-A^2}\sqrt{\left(\frac{\Delta r_\mathrm{tr}}{r_\mathrm{tr}}\right)^2 + \left(\frac{\Delta L}{L-h}\right)^2 + \left(\frac{\Delta h}{L-h}\right)^2}$$

we can simplify using trigonometric relations:

$$\frac{A}{1-A^2} = \frac{\sin\alpha}{1-\sin^2\alpha} = \frac{\sin\alpha}{\cos\alpha} = \tan\alpha$$

Draft

substituting into the error:

$$\Delta\alpha = \tan\alpha\sqrt{\left(\frac{\Delta r_{\text{tr}}}{r_{\text{tr}}}\right)^2 + \left(\frac{\Delta L}{L-h}\right)^2 + \left(\frac{\Delta h}{L-h}\right)^2}$$

after gathering the fractions with common denominators on the right:

$$\Delta\alpha = \tan\alpha\sqrt{\left(\frac{\Delta r_{\text{tr}}}{r_{\text{tr}}}\right)^2 + \frac{\Delta L^2 + \Delta h^2}{(L-h)^2}}$$

we can simplify further by replacing the distance errors with the respective $\sigma^{\text{top}}$,$\sigma^{\text{side}}$ errors (we note that it is possible to obtain smaller errors $L, h$ by measuring them physically. In our measurements we use image analysis for all the length measurements):

$$\boxed{\Delta\alpha = \tan\alpha\sqrt{\left(\frac{\sigma^{\text{top}}}{r_{\text{tr}}}\right)^2 + 2\left(\frac{\sigma^{\text{side}}}{L-h}\right)^2}} \qquad \textbf{(S.17)}$$

Finally we calculate the force error, from Eq. 2:

$$\boxed{\Delta F = \frac{1}{F}\sqrt{\begin{array}{l}\left(\frac{\Delta m}{m}\right)^2 + \left(\frac{\Delta L}{L}\right)^2 + \\ \left(\frac{\Delta l_{\text{c}}}{\sqrt{l_{\text{c}}}}\right)^2 + \left(\frac{\tan^2\alpha}{1+\tan^2\alpha}\Delta\alpha\right)^2\end{array}}} \qquad \textbf{(S.18)}$$

Where the error for $m$ is the scale accuracy, the error for $L$ is a distance error from the side view by Eq. S.13, and the error for $l_{\text{c}}$ is given by Eq. S.16

**Sensitivity estimation.** The maximal deflection for a given force occurs for contact near the lower tip of the rod, and $h$ is small - so $L, l_{\text{c}} \gg h$, If $h$ is small: $L \gg h$, we can ignore $h$ in the denominator, and approximate $l_{\text{c}} \approx L$. Assuming small deflection angles of the rod $\alpha \ll 1$, we get the approximate relations $\sin(\alpha), \tan(\alpha) \approx \alpha$. Substituting into Eq. 2:

$$F = \frac{mgL}{2l_{\text{c}}}\tan\alpha \ \ (\alpha \ll 1)$$
$$F \approx \frac{mgL}{2l_{\text{c}}}\alpha \ \ (l_{\text{c}} \approx L)$$
$$F_{\text{min}} \approx \frac{mg}{2}\alpha$$

**Minimal angle estimation.** To estimate the minimal detectable force in our system, we start with the resolution of the angle from the deflection by inverting Eq. 1:

$$\sin\alpha = \frac{r_{\text{tr}}}{L-h} \qquad \textbf{(S.19)}$$

Substituting the small angle approximation into Eq. S.19:

$$\alpha \approx \frac{r_{\text{tr}}}{L-h} \qquad \textbf{(S.20)}$$

Using the same approximations as above for $h$ and $\alpha$, we estimate the minimal detectable angle by taking the minimal deflection resolved in our system $r_{\text{tr}} = \sigma^{\text{top}}$ and substituting into Eq. S.20:

$$\boxed{\alpha_{\text{min}} \approx \frac{\sigma^{\text{top}}}{L}} \qquad \textbf{(S.21)}$$

Finally we estimate the minimal detectable force as the force obtained at the minimal resolvable angle according to Eq. S.21, which gives:

$$\boxed{F_{\text{min}} \approx \frac{mg}{2}\alpha_{\text{min}}} \qquad \textbf{(S.22)}$$

The approximate minimal angle in our setup is $\approx 0.005$ rad. Substituting into S.22 we get a minimal force of $\approx 400$ of the pendulum weight for our setup.

**Maximal force.** If we assume a maximal deflection angle of $\alpha_{\text{max}} = 15°$ ($\tan 15° \approx 1/4$), which is on the order of the maximal deflection in our setup ($\approx 12°$, see Fig. 6), and take a contact distance near the minimal contact distance in our experiments $\approx L/4$. We can make a rough estimation of the maximal detectable force for this system by substituting these values into Eq. 2 and get:

$$\boxed{F_{\text{max}} \approx \frac{mgL}{2\frac{L}{4}}\tan\alpha_{\text{max}} \approx \frac{mg}{2}} \qquad \textbf{(S.23)}$$

Such that the estimated maximal force for the system (Eq. S.23) will be of the same order of magnitude as the weight of the pendulum. This value will depend on the specific system parameters and the optical setup, but gives a useful ballpark estimation.

**Pendulum relaxation.** The moment of inertia of a rod of length L with constant density about the end point P is ((Landau, 1976)):

$$I_P = \frac{1}{3}mL^2 \qquad \textbf{(S.24)}$$

Draft

The angular frequency (or the maximal velocity?) of small oscillations for a physical pendulum with the hinge at the tip is approximately:

$$\omega_0 \simeq \sqrt{\frac{(L/2)mg}{I}} \quad \textbf{(S.25)}$$

and the period is:

$$T_{pend} = \frac{2\pi}{\omega_0} \simeq \sqrt{\frac{2L}{3g}} \quad \textbf{(S.26)}$$

,as shown below:

For a simple rod-pendulum at small angles approximately, the period is given by Morin (2008):

$$T_{pend} \approx 2\pi\sqrt{\frac{I_0}{mgr_{\oplus}}} \quad \textbf{(S.27)}$$

Where $I_0$ is the moment of inertia around the pivot point, $mg$ is the weight of the rod,$r_{\oplus}$ is the distance of between the hinge and the center of mass, $T_{pend}$ is the time of a full period of oscillation for the pendulum under gravity alone.

To simplify, we take an approximation that the rod mass is uniformly distributed (not hollow) - otherwise, a hollow cylinder has a somewhat complicated moment of inertia around the perpendicular axis.

The moment of inertia is given by:

$$I = \int r^2 dm$$

Where $r$ is the distance of the element $dm$ from the axis of rotation.

The moment of inertia for a uniformly distributed cylinder around its center for rotation about axis perpendicular to the long axis is given by:

$$I_{x,cm} = \frac{1}{4}mR^2 + \frac{1}{12}mL^2 \quad \textbf{(S.28)}$$

If we allow the radius to shrink the limit goes to the moment of inertia of a thin rod. We can obtain the moment of inertia for rotation about the rod tip by using the parallel axis (Huygens-Steiner) theorem:

$$I_p = I_{c.m} + md^2$$

Where the $I_{c.m}$ is the moment of inertia for the axis going through the center of mass from Eq. S.28, $m$ is the mass, $d$ is the distance of the axis from the center of mass. The moment of inertia around the rod tip is given by:

$$\begin{aligned} I_L &= \frac{1}{4}mR^2 + \frac{1}{12}mL^2 + m(L/2)^2 \\ &= \frac{1}{4}mR^2 + \frac{1}{3}mL^2 \end{aligned} \quad \textbf{(S.29)}$$

Where $I_L$ is the moment around the axis. Substituting $m, R, L$ for the typical values in our setup into Eq. S.27 yields a period of $T_{pend} \approx 1$ s, demonstrating the separation of timescales between the slow circumnutations and the fast mechanical relaxation (Eq. S.27). A quarter of this period can be taken as an approximate relaxation time between contact with the rod and stable contact.

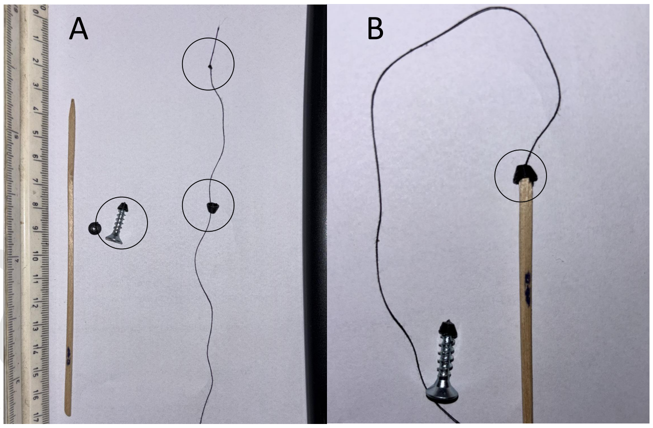


**Fig. S4. 3D printed caps for wooden rods.** (A) String with knot to prevent slipping, screw to enlarge hole and wooden rod. (B) Rod glued to cap, string above rod near cone tip.

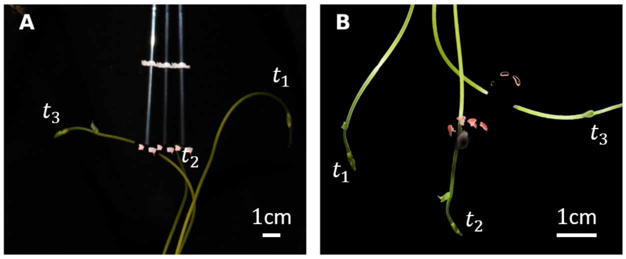


**Fig. S5. Overlay of top and side view images for impact of shoot on force measurement pendulum rod after t = 0, 15, 30 minutes.** (A) Side view. Tracked red point seen near lower tip of straw is moved over time with the motion of the stem. (B) Top view. The rod is barely visible at first, and over time a projection can be seen. Due to the short distance from the camera, large deflections will leave the frame (see stem at t=30 min), placing a practical limit on measured angles, though this is an adjustable property of the setup.

A small cone-cap was printed to match the rod tip. A small screw was used to increase hole size (as printing holes at that resolution proved difficult). A string was passed through the hole at the top of the cone, with a small knot preventing slipping out. The wooden rod was then glued to the cap. If the rod was too thin, the top of the rod was glued directly to the string.

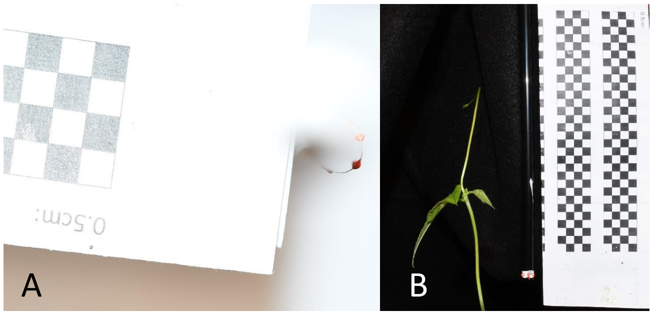


**Fig. S6. Pixel to cm conversion reference image.** (A) Top view. Diameter and square length are known sizes used for pixel conversion rates. Checkered page placed horizontally level with the bottom of rod. (B) Side view. Rod and square lengths used for pixel conversion rates. Checkered page placed at same plane as rod relative to side camera.

Physical length taken from images in pixels to get accurate pixel conversion rates, see Fig S6.

Draft
